# Dynamic regulation of translation quality control associated with ribosome stalling

**DOI:** 10.1101/2020.05.29.121954

**Authors:** Daniel H. Goldman, Nathan M. Livingston, Jonathan Movsik, Bin Wu, Rachel Green

## Abstract

Translation of problematic mRNA sequences induces ribosome stalling. Collided ribosomes at the stall site are recognized by cellular quality control machinery, resulting in dissociation of the ribosome from the mRNA and subsequent degradation of the nascent polypeptide and in some organisms, decay of the mRNA. However, the timing and regulation of these processes are unclear. We developed a SunTag-based reporter to monitor translation in real time on single mRNAs harboring difficult-to-translate poly(A) stretches. This reporter recapitulates previous findings in human cells that an internal poly(A) stretch reduces protein output ∼10-fold, while mRNA levels are relatively unaffected. Long-term imaging of translation indicates that poly(A)-containing mRNAs are robustly translated in the absence of detectable mRNA cleavage. However, quantification of ribosome density reveals a ∼3-fold increase in the number of ribosomes on poly(A)-containing mRNAs compared to a control mRNA, consistent with queues of many stalled ribosomes. Using single-molecule harringtonine runoff experiments, we observe the resolution of these queues in real-time by the cellular quality control machinery, and find that rescue is very slow compared to both elongation and termination. We propose that the very slow clearance of stalled ribosomes provides the basis for the cell to distinguish between transient and deleterious stalls, and that the human quality control apparatus predominantly targets the nascent protein rather than the mRNA.

## Main text

To maintain a functional proteome, the cell must faithfully express the genetic information encoded in DNA. Damaged or defective mRNAs, or other translation-related stresses, can stall the ribosome, generating generally toxic truncated protein products (*1*, *2*). The cell senses stalled ribosomes to initiate a quality control response involving ribosome rescue, nascent polypeptide degradation and in some cases, mRNA decay. Failure to resolve stalled ribosomes and degrade the resultant truncated proteins can induce proteotoxic stress and neurodegeneration (*3*–*5*).

A major class of mRNA that induces ribosome stalling and associated quality control includes those that lack a stop codon (*6*, *7*). Such mRNAs can be generated by premature cleavage and polyadenylation, a process that is enhanced during neuronal stimulation and in cancer cells (*8*, *9*). In the absence of a stop codon, the ribosome translates the poly(A) tail, where interactions with both the nascent poly-lysine peptide and poly(A) sequence cause the ribosome to stall (*10*–*12*). Stalled ribosomes are split from the mRNA, resulting in a 60S-peptidyl-tRNA complex (*13*–*15*). Subsequently, the E3 ligase Listerin ubiquitylates the nascent polypeptide, targeting it for proteasomal degradation (ribosome-associated quality control, or RQC) (*5*, *16*). In yeast, translation stalling triggers mRNA decay, proceeding through the actions of the canonical Xrn1 exonuclease and the quality control-specific endonuclease Cue2 (*17*, *18*). Experiments exploring whether related mRNA decay occurs in mammalian cell lines have produced mixed results (*7*, *19*, *20*).

Recent work established that initiation of the quality control response depends on ribosome collisions at the stall site (*21*), which are recognized by binding of the ubiquitin ligase ZNF598 to the collided di-ribosome (*22*–*25*). However, given the density of translating ribosomes and the abundance of even modest pause-inducing motifs, as many as 10% of ribosomes on highly-translated mRNAs are involved in collisions (*26*, *27*). It is unclear how the cell distinguishes between regulatory or routine pauses and those involving problematic mRNAs that should be targeted for quality control. Although stalling is inherently a kinetically-defined process, there have been no *in vivo* kinetic measurements of stalling. Thus, it is not known how long ribosomes spend in a paused state before being removed from the mRNA. Additionally, it is unclear whether RQC is coupled with mRNA decay in human cells, or how ZNF598 contributes to ribosome rescue. We set out to address these fundamental questions through real-time observation of ribosome stalling and ensuing quality control, observing the dynamics of this process in living cells for the first time.

To monitor translation quality control on single mRNAs in human cells, we implemented the SunTag method (*28*–*32*). In this system, a reporter mRNA encodes tandem repeats of the SunTag epitope near the 5’ end of the open reading frame (Figure 1A). Upon translation of each SunTag epitope, a single chain variable fragment (scFV) of a GCN4 antibody fused to super folder GFP (scFV-sfGFP) binds the nascent polypeptide, reporting on translational activity. The mRNA is labeled red and tethered to the cell membrane through binding of MS2 coat-binding protein fused to Halotag and a CAAX motif (MCP-Halo-CAAX). It was previously demonstrated that tethering the mRNA limits diffusion of the reporter, allowing stable imaging without affecting translational output (*30*). Downstream of the SunTag array, Nano Luciferase (NLuc) allows for measurement of bulk protein output, while an auxin-inducible degron (AID) enables controlled depletion of the fully-synthesized polypeptide to reduce fluorescence background (*28*, *33*).

**Fig.1.**
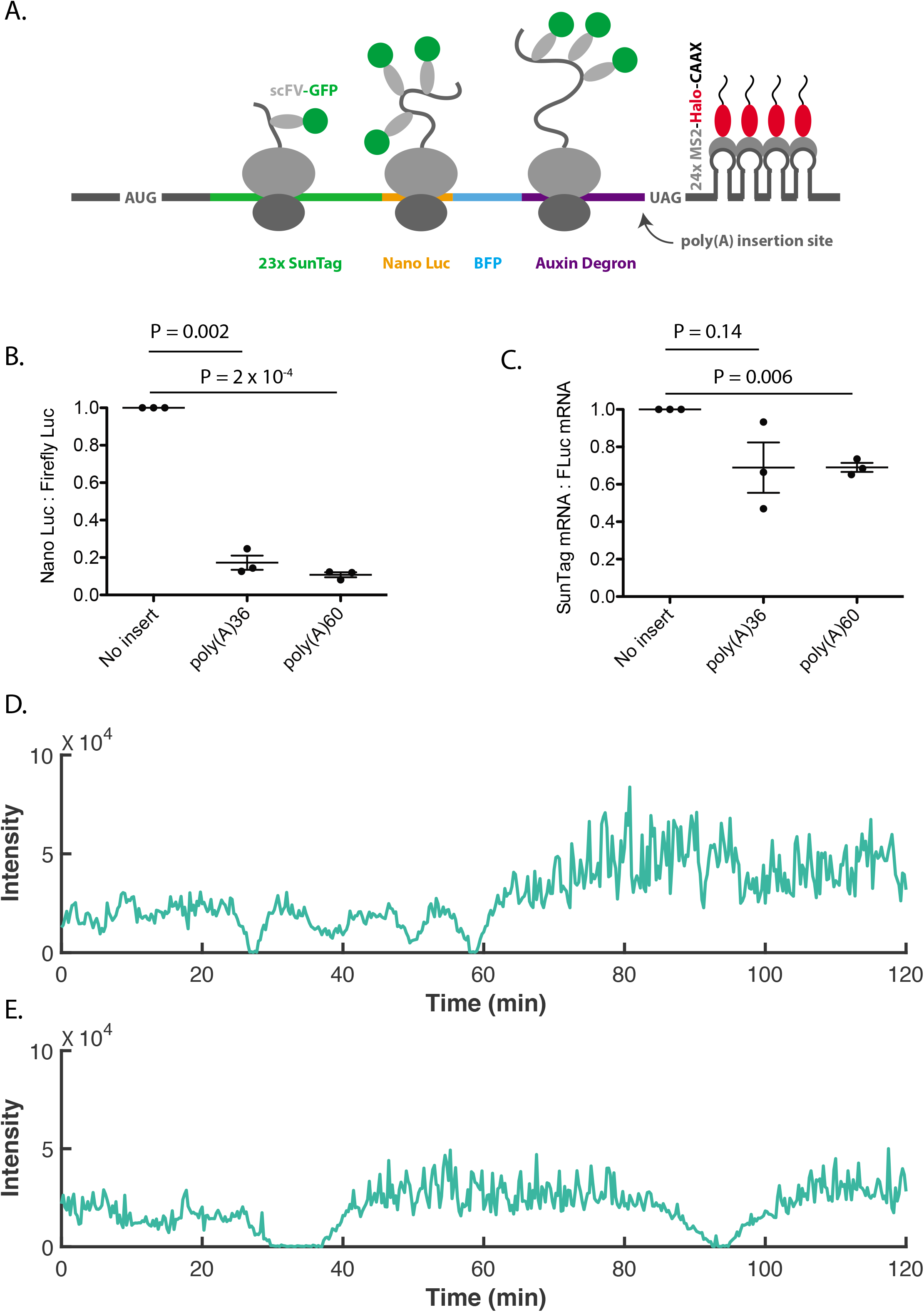
SunTag reporters for monitoring translation quality control on single mRNAs. (A) Reporter schematic depicting ribosomes with scFV-GFP-bound nascent polypeptides. The open reading frame of the reporter contains 1253 sense codons. (B) Luciferase assay to measure reporter protein output. NLuc signal is normalized to Firefly Luciferase signal from a cotransfected plasmid and the resulting ratios are normalized to the no insert reporter. Error bars represent the sem of 3 independent experiments. p-values calculated by paired-sample t-test. (C) Quantification of mRNA levels by RT-qPCR. SunTag mRNA levels are normalized to mRNA levels of the co-transfected Firefly Luciferase plasmid and the resulting ratios are normalized to the no insert reporter. Error bars represent the sem of 3 independent experiments. p-values calculated by paired-sample t-test. (D) Example trace of SunTag intensity over time for a no insert mRNA. (E) Example trace of SunTag intensity over time for a poly(A)60 mRNA.

We generated versions of the SunTag reporter without a stalling sequence (“no insert”) or including a stretch of either 36 or 60 adenosines shortly upstream of stop codons in all three frames (“poly(A)36” or “poly(A)60”). Direct mRNA sequencing using Oxford Nanopore technology revealed some cryptic splicing in the repetitive SunTag region of the reporters; importantly, the features of the most abundant mis-spliced isoforms are incompatible with expression of both NLuc and SunTag, rendering their translation products undetectable by luciferase assay and imaging (Figure S1). Nevertheless, our data highlight the importance of assessing the integrity of SunTag mRNAs by sequencing. Upon transfection of the reporters into U-2OS cells, luciferase assays revealed that total protein output is ∼6- or ∼9-fold suppressed on poly(A)36 or poly(A)60, respectively, relative to the no insert reporter (Figure 1B). By comparison, mRNA levels of the poly(A) reporters are reduced by ∼30%, suggesting only a minor contribution from mRNA loss (Figure 1C). These observations are consistent with previous work, in which ribosome stalling on poly(A)60 induced nascent polypeptide degradation and a comparable reduction in reporter expression (*19*).

Although our measurements of protein and mRNA levels are consistent with ribosome stalling leading primarily to ribosome rescue and nascent polypeptide degradation, the data could also be explained by reduced efficiency of translation initiation on the poly(A) reporters. To determine whether this is the case, we introduced a P2A sequence upstream of the poly(A) insertion site (Figure S2A). Translation of P2A causes the ribosome to release the peptide at an intermediate point during ribosome elongation, thus dissociating NLuc from the ribosome prior to reaching the stalling site and insulating NLuc from the effects of RQC (*34*). If stalled ribosomes on poly(A) trigger RQC, insertion of the P2A sequence should restore protein levels; if the data reflect reduced translation initiation efficiency, the P2A sequence should have no effect. Insertion of the P2A sequence resulted in near complete restoration of protein output from the poly(A)60 reporter (to ∼80% of no insert levels), indicating that degradation of the nascent polypeptide coincident with stalling is responsible for most of the suppressed protein output (Figure S2B). Additionally, experiments in HEK293T cells that do not express the scFV-sfGFP or MCP-Halo-CAAX proteins recapitulated those in U2OS cells, demonstrating that protein output is not affected by mRNA tethering, antibody binding to the nascent peptide or the specific cell line (Figure S3). Together, the luciferase and RT-qPCR data indicate that the majority of ribosomes stall on poly(A), resulting in ribosome rescue and nascent polypeptide degradation.

To monitor the translation status of mRNAs in real-time, we transfected the no insert and poly(A)60 SunTag reporters into cells and imaged steady-state translation for 3 h, observing mRNAs for a median time of ∼40-45 min (Figure S4, movies S1 and S2). Translation on both the no insert and poly(A)60 reporters exhibits a characteristic bursting pattern, in which mRNAs cycle between a translationally active and inactive state (Figure 1D-E) (*28*, *30*, *32*). The no insert and poly(A) mRNAs are similarly active, found in the “on” state 81% of the time. Translating mRNAs are generally observed in one of three phases: ramp-up, in which a previously inactive mRNA is being loaded with ribosomes; steady-state translation, in which the ribosome load remains roughly constant over time; ramp-down, in which the ribosome load is gradually reduced until complete translation shut-down. Importantly, removal of a ribosome during normal termination or by rescue on poly(A) both result in loss of one SunTag nascent polypeptide and thus a reduction in green fluorescence intensity at the translation site. The observation of both ramp-ups and ramp-downs indicates that both reporters are actively translated, undergoing translation initiation, elongation, and ribosome removal (by either normal termination or ribosomes rescue).

A recent study employing the SunTag method estimated that an average of 8 ribosomes translate a premature termination codon (PTC)-containing mRNA before the transcript is cleaved by the endonuclease SMG6 and targeted for nonsense-mediated decay (NMD) (*35*). In this case, mRNA cleavage was visualized as an abrupt separation of red and green signals (foci separation) as SMG6 cleaved the mRNA at the terminating ribosome and severed upstream translating ribosomes from red signal localized to the 3’UTR stem loops. Additionally, previous studies have reported similar endonucleolytic cleavage of mRNA associated with translation stalling in yeast (*17*, *18*). Given that the poly(A) sequence in our reporter is located at the 3’ end of a long open reading frame (ORF; 1275 codons), a cleavage at or near the poly(A) sequence would sever most translating ribosomes from the MS2 and PP7 stem loops that label the mRNA and tether it to the membrane (Figure S5). We thus reasoned that if cleavage were taking place, we might be able to observe a similar abrupt separation of red signal from green signal on a translating mRNA (“foci separation”). Although we are able to observe foci separation, it is a rare event, occurring with similar frequency for the no insert and poly(A)60 reporters (0.018 h^−1^ or 0.024 h^−1^, or once in 5.5 h or 4.2 h of total translation time, respectively) (movie S3). These data are consistent with the low level of background cleavage observed on a non-PTC-containing control reporter in the NMD study. Thus, we do not detect specific endonucleolytic cleavage of the poly(A)60 reporter. Taken together with the luciferase and RT-qPCR measurements, these data suggest that the poly(A)60 reporter is robustly translated and subject to nascent polypeptide degradation but little to no mRNA decay. We thus focused on the fate of ribosomes translating these mRNAs.

Although it is widely accepted that ribosomes stall on poly(A), the dynamics of this process have never been directly observed. For example, neither the duration of a stall, nor the buildup (number of accumulated ribosomes) caused by a stall have been measured. We first set out to quantify the ribosome load on each SunTag reporter. To control the amount of cell to cell variation in mRNA expression, we generated cell lines for stable expression of the SunTag reporters from a single locus; protein output from these cell lines recapitulated the patterns observed with the transfected reporters characterized above (Figure S6). We next used single-molecule fluorescence in-situ hybridization for detection of individual mRNAs (smFISH) combined with immunofluorescence (IF) detection of GFP to quantify the SunTag signal (smFISH-IF) (*36*). Bright green spots in the IF channel colocalized with red spots in the smFISH channel, reflecting translating mRNAs, while un-colocalized dim green spots in the IF channel represent single fully synthesized polypeptides released from the ribosome and bound by scFV-GFP (Figure 2A) (*28*). Although these single polypeptides are rapidly degraded owing to the AID sequence, they are stable enough for some of them to be detected by fixed-cell imaging.

**Fig.2.**
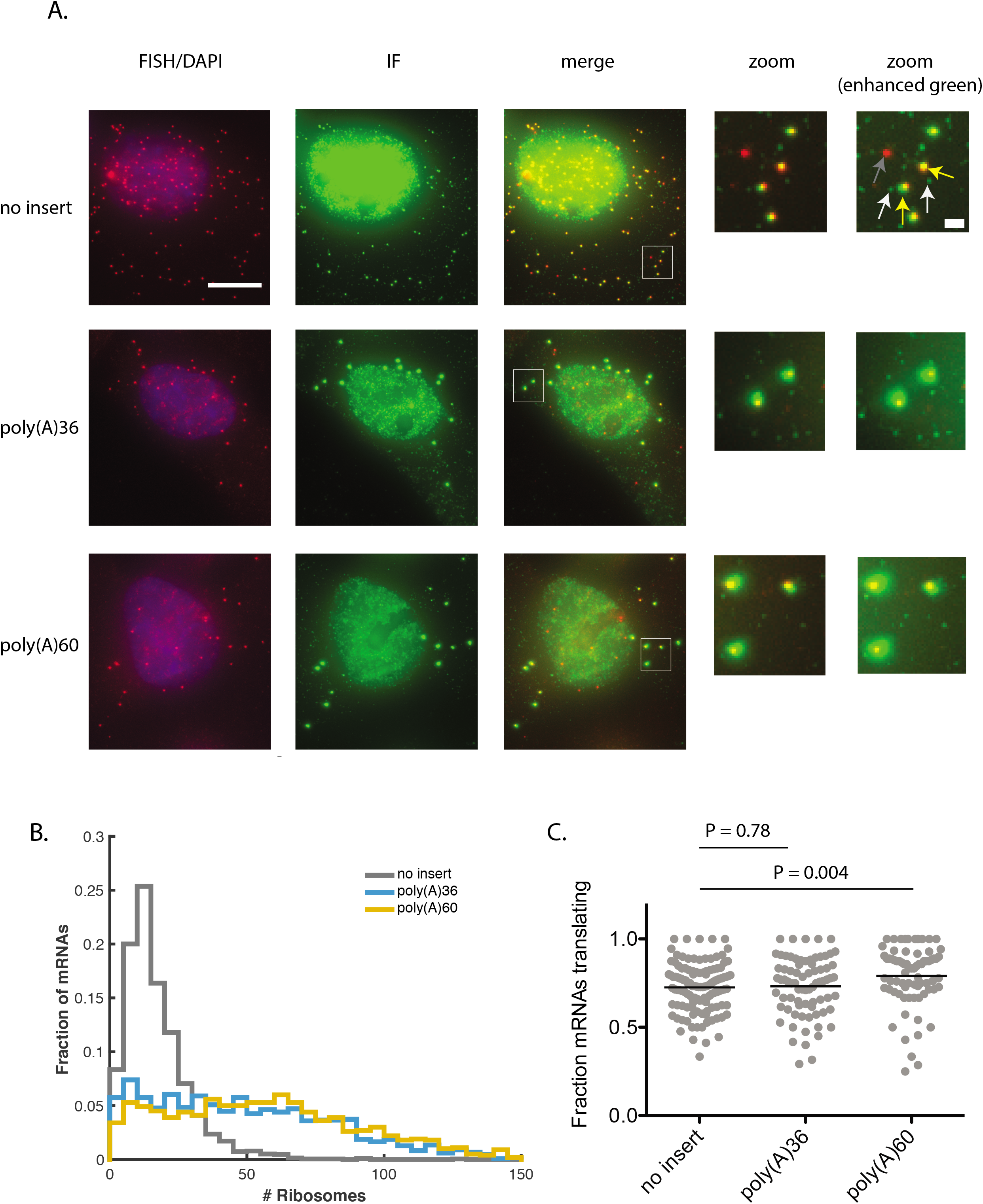
Poly(A) reporters have increased ribosome load relative to the no insert reporter. (A) Example cells from smFISH-IF experiment. Red spots that do not colocalize with green reflect un-translating mRNAs (gray arrow on the right-most panel), red spots that colocalize with green reflect translating mRNAs (yellow arrows), and dim green spots that do not colocalize with red reflect fully-synthesized and released polypeptides (white arrows). Spot intensity is directly comparable among these images, indicating that poly(A) reporters are brighter in the green channel and thus comprise more translating ribosomes than the no insert reporter. White boxes drawn on “merge” images indicate zoom regions for right-most panels. Scale bar in top left image: 10 microns. Scale bar in top “enhanced zoom” image: 1 micron. (B) Quantification of number of ribosomes per mRNA. Data compiled from two independent experiments. 110-134 cells, 1020-2169 mRNAs per condition. p-value for comparison of no insert to poly(A)36: < 1 × 10^−100^; p-value for comparison of no insert to poly(A)60: < 1 × 10^−100^. p-values calculated by two-sample t-test after correction of distribution skewness (see supplemental text for details). (C) Fraction of mRNAs actively translating (calculated only for cells with more than 5 mRNAs). Each point represents one cell, black lines indicate mean. p-values calculated by two-sample t-test.

Importantly, quantification of the intensity of single polypeptides revealed no difference in brightness among the three samples (Figure S7). However, the SunTag intensity co-localized with poly(A)36 and poly(A)60 mRNAs appeared substantially brighter in the IF channel than the no insert mRNAs (Figure 2A). We determined the mean integrated intensity of the released single polypeptides and used this value to estimate the number of translating ribosomes on each mRNA (supplementary text). While the no insert reporter was occupied by an average of 17 ribosomes, the poly(A)36 and poly(A)60 reporters were occupied by an average of 50 and 58 ribosomes, respectively (Figure 2B, S8); all three reporters exhibited a similar fraction of mRNAs in a translating state (Figure 2C). The average ribosome load on the no insert reporter reflects a density of one ribosome every ∼75 codons, approximately 7.5-fold below the maximum packing density assuming a ribosome footprint of 10 codons (*37*). In contrast, the average density on the poly(A)60 reporter is almost half maximal, indicating queues of many collided ribosomes that extend well upstream of the poly(A) sequence. The slight decrease in ribosome load on the poly(A)36 reporter relative to the poly(A)60 reporter is consistent with the respective increase in luciferase output (Figure 1B) and indicates that slightly more ribosomes are able to read through the shorter poly(A)36 sequence and terminate at the stop codon. Overall, the accumulation of ribosomes on the poly(A) reporters suggests that rescue of stalled ribosomes is slow compared to normal termination at a stop codon. However, it is not possible to ascertain the relative removal rate of stalled ribosomes from these static images.

If ribosome rescue is indeed slow compared to normal termination, the total residence time of a ribosome should be longer on the poly(A) reporter than on the no insert reporter. To determine if this is the case, we monitored translation ramp-down events observed during live-cell imaging (for examples, see Figure 1D, starting at ∼25 min and ∼55 min; also, see Figure 1E, starting at ∼25 min and ∼85 min). During these ramp-down events, mRNAs initially being translated at steadystate are cleared of ribosomes, either by elongation and termination (for the no insert reporter), or by elongation and ribosome rescue (for the poly(A) reporters). If rescue of ribosomes stalled at poly(A) is slow, translational ramp-down should take longer on poly(A)60 mRNAs than on no insert mRNAs. Indeed, we find that translation ramp-down on poly(A)60 mRNAs is slower than on no insert mRNAs, as measured by the time between steady-state translation (the plateau) and complete shut-down (Figure S9). These data suggest that ribosome clearance from the poly(A)60 reporter is limited by the rate of ribosome rescue; however, this measurement is complicated by the fact that these mRNAs are not synchronized with respect to translational shut-down.

To more thoroughly investigate the timing of ribosome rescue, we performed live-cell imaging after addition of the drug harringtonine, which permits initiation but blocks formation of the first peptide bond during translation (*38*). Upon treatment with harringtonine, all newly initiated ribosomes are prevented from elongating through the mRNA (remaining bound at the initiation site), while actively elongating ribosomes are able to finish translating (*39*). This strategy allows us to synchronize mRNA molecules at the time of drug application and observe ribosome clearance in real time. In general, treatment with harringtonine resulted in a gradual loss of green fluorescence (Figure 3A, movies S4-S6). Inspection of individual molecules reveals that loss of signal proceeds more slowly on the poly(A) reporters than on the no insert reporter (Figure 3A-C). This delay is consistent with the slower translational ramp-down observed on poly(A)60 during steady-state (no drug) experiments (Figure S9).

**Fig.3.**
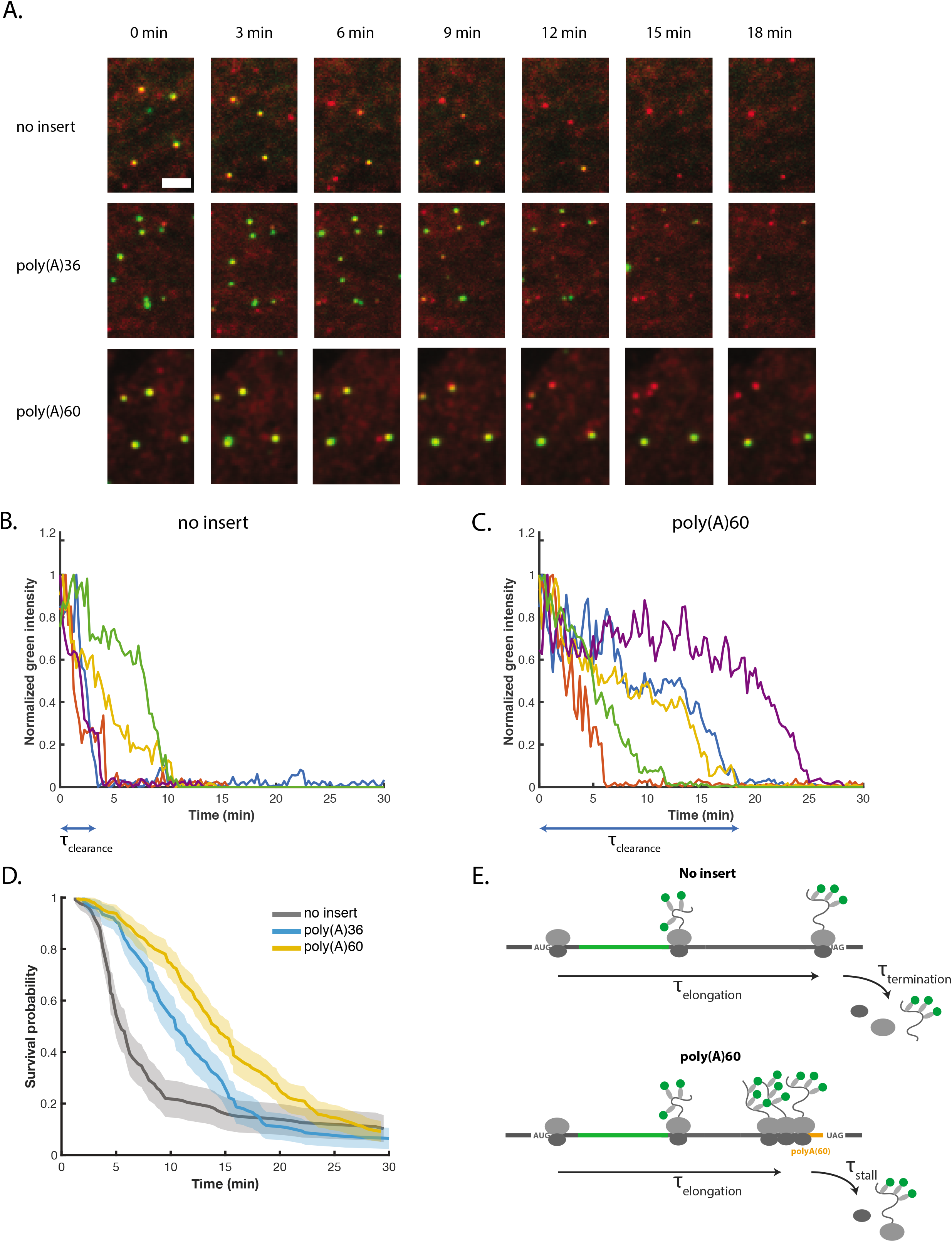
Ribosome clearance is delayed on poly(A) mRNAs. (A) Snapshots from movies at indicated post-harringtonine timepoints. Green spot intensity is directly comparable among these images. Scale bar in top left image: 3 microns. (B) Example traces of green intensity over time for single no insert mRNAs post-harringtonine treatment. The signal for each molecule is normalized to its maximum intensity. Blue double arrow below the x-axis indicates *τ_clearance_* for the blue example trace. (C) Example traces of green intensity over time for single poly(A)60 mRNAs post-harringtonine treatment. The signal for each molecule is normalized to its maximum intensity. Blue double arrow below the x-axis indicates *τ_clearance_* for the blue example trace. (D) Cumulative survival probability of green signal on mRNAs post-harringtonine treatment. Shaded area represents 95% confidence bounds computed using Greenwood’s formula. 8-11 cells, 127-199 mRNAs per condition. (E) Cartoon illustrating the kinetic steps comprising *τ_clearance, no insert_* (top) and *τ*_*clearance,poly*(*A*)_ (bottom). *τ_clearance, no insert_* = *τ_elongation_* + *τ_termination_*, while *τ*_*clearance*,*poly*(*A*)_ = *τ_elongation_* + *τ_stall_*.

For each mRNA, we measured the time between harringtonine addition and complete loss of green fluorescence (*τ_clearance_*), corresponding to removal of the final ribosome (for examples of *τ_clearance_* determination, see blue double arrows in Figures 3B and 3C). To evaluate the population of clearance times, we generated a cumulative survival distribution of *τ_clearance_* (Figure 3D). For the no insert reporter, survival probability drops sharply around a median time of ∼5.5 min, consistent with the expected time for a ribosome near the 5’-end of the mRNA to elongate an ORF of this length (1253 codons at 3.5-5.6 codons/s is 3.7-6.0 min) (*28*, *30*, *39*). By comparison, loss of green signal is markedly delayed for the poly(A)36 and poly(A)60 reporters, with a median clearance time ∼5 and ∼8 min longer than the no insert reporter, respectively. Faster clearance of poly(A)36 relative to poly(A)60 suggests that ribosomes more often read through the shorter poly(A)36 sequence, increasing the effective rate of ribosome removal; this interpretation is consistent with luciferase data indicating higher protein output on poly(A)36 (Figure 1B). The distribution of *τ_clearance_* for all reporters reflects the stochasticity of underlying processes, including for example the location of the 5’-most ribosome. As seen from the non-zero baseline of the survival curves, ∼10% of mRNAs do not clear within the 30 min experimental time window for both no insert and poly(A) constructs; although we are unsure why these mRNAs fail to clear, this observation is consistent with previous reports (*30*).

To estimate the time a ribosome spends in the queue before being removed from the mRNA, we considered the difference in *τ_clearance_* between the no insert and poly(A) reporters. For the no insert reporter, *τ_clearance_* reflects the time for the 5’-most ribosome to elongate and terminate at the stop codon (*τ_clearance, no insert_* = *τ_elongation_* + *τ_termination_*). For the poly(A) reporters, we assume that the delay in *τ_clearance_* reflects the time required to rescue stalled ribosomes from the queue and not for ribosomes to read-through the poly(A) sequence. This assumption is supported by our luciferase data, indicating that the majority of ribosomes on the poly(A) reporters are rescued from the mRNA, and their nascent peptides degraded, before reaching the stop codon (Figure 1B). Thus, for the poly(A) reporters, *τ*_*clearance,poly*(*A*)_ = *τ_elongation_* + *τ_stall_* is the time to rescue the last stalled ribosome in the queue (Figure 3E).

In our estimate, we also assume that the time for a ribosome to terminate at the stop codon is small compared to the total time required to elongate the 1253-codon ORF (*τ_termination_* ≪ *τ_elongation_* and thus *τ_clearance,no insert_* ≈ *τ_elongation_*). This assumption is supported by relative peak heights of ribosome footprinting data at stop codons compared to sense codons, which indicates that the termination rate is at most 10-fold slower than the time to elongate an average codon (*39*, *40*). Thus, *τ_termination_* is on the order of seconds. Because the ORF upstream of poly(A) is identical to the ORF of the no insert reporter (both 1253 codons), *τ*_*clearance,poly*(*A*)_ ≈ *τ_clearance, no insert_* + *τ_stall_* Thus *τ_stall_* ≈ *τ*_*clearance,poly*(*A*)_ – *τ_clearance,no insert_*. Using median values for this calculation, *τ_stall_* ≈ 5 min for poly(A)36 and *τ_stall_* ≈ 8 min for poly(A)60. Since no new ribosomes translate the SunTag sequence after harringtonine treatment, *τ_stall_* reflects the average time that the final ribosome spends in the queue before leaving the mRNA, which is on the order of minutes. The observation that ribosome rescue is very slow compared to both normal elongation and termination provides a basis for the cell to distinguish between innocuous and deleterious collisions: Transient collisions involving actively elongating ribosomes will likely resolve within seconds, evading detection by the quality control machinery, while persistent collisions can be targeted by the quality control machinery.

In light of the well-established role of the E3 ligase ZNF598 in processing stalled ribosomes and targeting nascent peptides for decay, we were interested in assessing its impact in our system. A previous study demonstrated that depletion of ZNF598 *in vivo* increased the ability of ribosomes to read through a poly(A) sequence and synthesize a downstream reporter, highlighting its role in detection of stalled ribosomes (*19*). However, it is unclear mechanistically how ZNF598 activity limits readthrough of poly(A). In one scenario, ZNF598 accelerates splitting of stalled ribosomes from the mRNA (i.e. through recruitment of the helicase-like protein ASCC3 (*41*)) and thus limits readthrough; alternatively, ZNF598 simply stabilizes collided ribosomes and thereby limits readthgrough. These two models predict different outcomes for measurements of ribosome residence time at the stall site: if ZNF598 accelerates splitting, then residence time should be shortened, while if ZNF598 simply stabilizes the stall, then residence time should be lengthened.

In line with previous work, depletion of ZNF598 by siRNAs (Figure S10) in our system resulted in a small but reproducible increase in NLuc signal from the poly(A) reporter relative to the no insert reporter, indicating that ribosomes are indeed more likely to read through the poly(A) stretch and terminate at the stop codon (Figure 4A)(*19*). To determine how ZNF598 affects the residence time of stalled ribosomes, we performed harringtonine run-off experiments in cells depleted of ZNF598. While translation of the no insert reporter was unaffected by ZNF598 knockdown, *τ_clearance_* for the poly(A) reporter increased by an additional ∼14 min. (Figure 4B, movies S7-S10). Given the observed delay in ribosome clearance, we predicted that ZNF598 depletion should also cause an increased accumulation of stalled ribosomes on the poly(A)60 reporter. Consistent with this prediction, we observed a modest increase in ribosome load on poly(A)60 under ZNF598 depletion conditions, while ribosome load was insensitive to ZNF598 levels on the no insert reporter (Figure 4C and 4D, S11). Given that the luciferase assay indicated only a slight increase in the number of ribosomes reading through the poly(A) sequence upon ZNF598 knockdown (Figure 4A; most ribosomes are still rescued before reaching the stop codon), the clearance delay observed by harringtonine run-off reflects a delay in removal of stalled ribosomes (as opposed to the time to read through the poly(A) sequence). Thus, ZNF598 contributes to quality control by accelerating splitting of stalled ribosomes from the mRNA.

**Fig.4.**
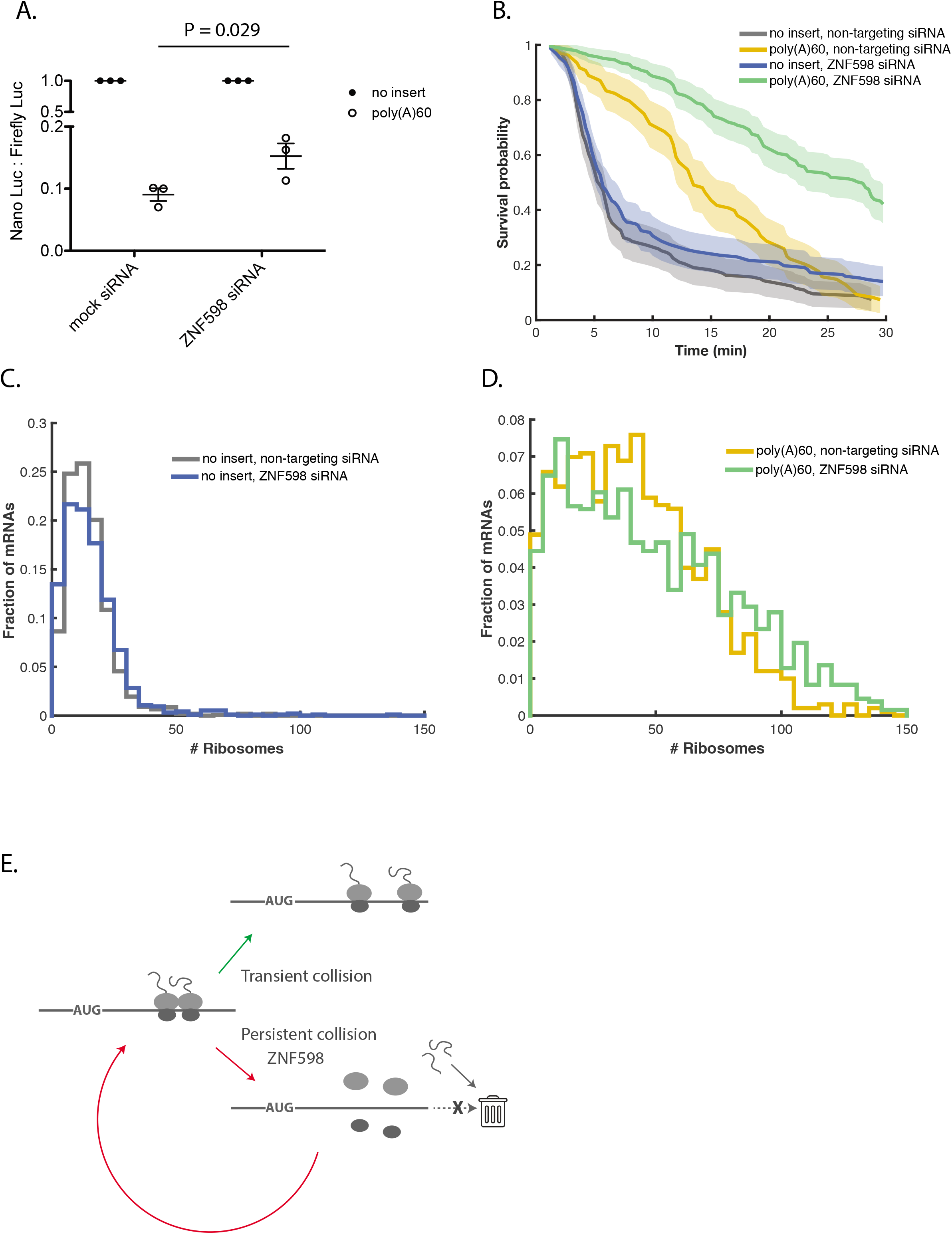
ZNF598 knock-down further delays rescue of stalled ribosomes. (A) Luciferase assay to measure reporter protein output under ZNF598 depletion conditions. NLuc signal is normalized to Firefly Luciferase signal from a co-transfected plasmid and the resulting ratio is normalized to the no insert reporter within each siRNA condition. Error bars represent the sem of 3 independent experiments. p-values calculated by paired-sample t-test. (B) Cumulative survival probability for harringtonine runoff experiment under ZNF598 depletion conditions. Shaded area represents 95% confidence bounds computed using Greenwood’s formula. 7 cells, 108-185 mRNAs per condition. (C and D) FISH-IF measurement of number of ribosomes on (C) no insert reporter under non-targeting siRNA (mean: 15 ribosomes) or ZNF598-targeting siRNA (mean: 14 ribosomes) conditions (D) poly(A)60 reporter under non-targeting siRNA (mean: 42 ribosomes) or ZNF598-targeting siRNA (mean: 50 ribosomes) conditions. Data compiled from two independent experiments. 82-124 cells, 951-1540 mRNAs per condition. p-value for comparison of no insert, nontargeting siRNA to no insert, ZNF598 siRNA: 0.25; p-value for comparison of poly(A)60, non-targeting siRNA to poly(A)60 ZNF598 siRNA: 2.1 × 10^−6^. p-values calculated by two-sample t-test after correction of distribution skewness (see supplemental text for details). (D) Model for how the cell senses and responds to translation pauses. Transient collisions are resolved when the lead ribosome resumes translating on the timescale of normal elongation. When a deleterious stall induces a long-lived collision, recognition by ZNF598 triggers ribosome rescue and nascent polypeptide degradation. mRNA decay is only weakly coupled or totally uncoupled from ribosome rescue, allowing ongoing translation and ribosome rescue to occur on stall-inducing mRNAs.

Altogether, our work reveals the temporal regulation of translation quality control in response to ribosome collisions. Rescue of collided ribosomes is slow compared to elongation, explaining how transient collisions can resolve before they are targeted for quality control, while ZNF598 specifically targets prolonged collisions (Figure 4E). We observe steady-state translation over minutes to hours in the absence of mRNA decay, suggesting that problematic mRNAs undergo many rounds of translation and ongoing ribosome rescue over their lifetime. This finding is contrary to the yeast system, in which mRNA decay is a robust feature of the quality control response to translation stalling (*7*, *17*). Additionally, our observations are in stark contrast with NMD in a human system, in which mRNA cleavage occurs rapidly after an average of only eight ribosome encounters(*35*). It is curious that the quality control response to translation stalling in human cells favors ribosome rescue and polypeptide degradation over mRNA decay. Our singlemolecule approach lays the groundwork for future studies on the regulation of translation quality control and the role of a rapidly expanding cast of cellular quality control factors.

## Supporting information

Oligonucleotide Sequences

## Acknowledgements

We thank the Johns Hopkins University Genetic Resources Core Facility for sequencing services. R.G. is an investigator of the Howard Hughes Medical Institute. D.H.G. is a Damon Runyon Fellow supported by the Damon Runyon Cancer Research Foundation (DRG-2280-16). N.M.L. is supported by NIH Training Grant T32 GM007445. We thank Boris Zinshteyn for critical discussions and feedback throughout the duration of the project.

## Supplementary Materials

Materials and Methods

Figures S1-S11

Movies S1-S10

References (*1-10*)

