## Supplementary Information for "Dynamic regulation of translation quality control associated with ribosome stalling"

**This PDF file includes:**

Materials and Methods
Supplementary Text
Figs. S1 to S11
Captions for Movies S1 to S10

**Materials and Methods**

Plasmids

The reporters used in this study are derived from that of an initial study demonstrating the SunTag method for monitoring translation, with several modifications (*1*). First, reporters were cloned into a pcDNA5 vector, and thus are expressed from a CMV promoter, instead of a UbC promoter. Additionally, a Nano Luciferase was inserted downstream of the SunTag array, and most of the b-actin 3'UTR was removed. The original SunTag array comprises 24 repeats; however, we inactivated the second repeat (g to c substitution, swapping valine for alanine) in an effort to mitigate cryptic splicing. The resulting plasmid, pcDNA_CMV_ST (1253 codons), was used to generate the poly(A) reporters: pcDNA_CMV_ST_A36 and pcDNA_CMV_ST_A60. These plasmids differ from pcDNA_CMV_ST by a stretch of 36 or 60 adenosines, respectively, plus two additional codons (coding for alanine and serine) inserted two codons upstream of the stop codon. If frameshifting occurs on the poly(A) sequence, a stop codon is encountered either 4 nt upstream or downstream of the in-frame stop codon. In order to generate MCP-Halo-CAAX, MCP-tagRFP-CAAX was first made by inserting the CAAX motif between the AgeI and ClaI sites of pubc-

2xMCP-tagRFP. The CAAX motif was removed by PCR from pHR-PP7-2xmCherry-CAAX (Addgene #74925). The final MCP-Halo-CAAX plasmid was generated by inserting a HaloTag between the AgeI and XbaI cut sites.

Cell culture

Cells were cultured in DMEM (Thermo Fisher, #11995) supplemented with 10% FBS (Thermo Fisher, #A3160401) at 37°C and 5% $CO_2$. For smFISH-IF experiments, cells were cultured in the presence of 100 units penicillin and 0.1 mg/mL streptomycin (Sigma Aldrich, P0781). For steady-state live-cell imaging experiments (in the absence of harringtonine), cells were cultured beforehand in the presence of pen/strep, but pen/strep was not present during imaging. For all other experiments, cells were cultured (and experiments performed) in the absence of pen/strep.

Luciferase assays

For experiments in which SunTag reporters were transfected, U-2 OS cells stably expressing MCP-Halo-CAAX and scFV-sfGFP were seeded in a 96-well plate at a density of 7000 cells per well. The following day, wells were transfected with 29 ng of reporter plasmid and 2 ng of firefly luciferase-expressing normalizer plasmid (pcDNA5_FRT_TO_EGFP_AID_Luciferase, a gift from Dr. Andrew Holland) using X-tremeGENE 9 DNA Transfection Reagent (Roche, #06365787001), at a ratio of 5 ul reagent to 1 ug DNA to 125 ul Opti-MEM Reduced Serum Medium (Thermo Fisher, #51985034). Approximately 24 h later, Nano and Firefly Luciferase activities were measured using the NanoGlo Dual-Luciferase Reporter Assay System (Promega, #N1630) in a Synergy H1 microplate reader (BioTek). Nano Luciferase values for either two or three technical replicate wells were averaged, as were the Firefly Luciferase values for the same wells. The ratio of Nano Luciferase to Firefly Luciferase was then taken, and the ratio for each reporter was normalized to that of the no insert reporter. Each experiment was performed three times and the average and standard error of the mean reported.

For luciferase assays performed using stable cell lines, U-2 OS cells stably expressing scFV-sfGFP and harboring a genomic copy of the SunTag reporters were seeded in a 96-well plate at a density of 7000 cells per well. The following day, wells wells for cells harboring each reporter were induced by adding doxycycline hyclate (Millipore Sigma, #D9891) to a final concentration of 2 ug/ml. Approximately 24 h later, Nano Luciferase activity was measured using the NanoGlo Dual-Luciferase Reporter Assay System (Promega, #N1630) in a Synergy H1 microplate reader (BioTek). Nano Luciferase values for either two or three technical replicates were averaged and the resulting value normalized to that of the no insert reporter. Each experiment was performed three times and the average and standard error of the mean reported.

For luciferase experiments performed under ZNF598 depletion, cells were seeded in a 96-well plate at a density of 7000 cells per well. The following day, cells were transfected with a pool of 4 uM ZNF598-targeting siRNAs (Horizon, #L-007104-00-0005) or a pool of 4 uM non-targeting siRNA (Horizon, #D-001810-10-05) using RNAiMax (Thermo Fisher, #13778) according to manufacturer protocol. Approximately 24 h later, cells were transfected with a second dose siRNAs. Approximately 4 h later, the media was changed to fresh DMEM + 10% FBS and triplicate wells were transfected with 29 ng of reporter plasmid and 2 ng of firefly luciferase-expressing normalizer plasmid (pcDNA5_FRT_TO_EGFP_AID_Luciferase, a gift from Dr. Andrew Holland) using X-tremeGENE 9 DNA Transfection Reagent (Roche, #06365787001), at a ratio of 5 ul reagent to 1 ug DNA. Approximately 24 h later, Nano and Firefly Luciferase activities were measured using the NanoGlo Dual-Luciferase Reporter Assay System (Promega,

#N1630) in a Synergy H1 microplate reader (BioTek). Nano Luciferase values for either two or three technical replicate wells were averaged, as were the Firefly Luciferase values for the same wells. The ratio of Nano Luciferase to Firefly Luciferase was then taken, and the ratio for each reporter was normalized to that of the no insert reporter. Each experiment was performed three times and the average and standard error of the mean reported.

Quantifying reporter mRNA levels

U-2 OS cells stably expressing MCP-Halo-CAAX and scFV-sfGFP were seeded in a 12-well plate at a density of 60,000 cells per well. The following day, each well was transfected with 290 ng of reporter plasmid and 20 ng of firefly luciferase-expressing normalizer plasmid (pcDNA5_FRT_TO_EGFP_AID_Luciferase, a gift from Dr. Andrew Holland) using X-tremeGENE 9 DNA Transfection Reagent (Roche, #06365787001), at a ratio of 5 ul reagent to 1 ug DNA to 125 ul Opti-MEM Reduced Serum Medium (Thermo Fisher, #51985034). Approximately 24 h later, cells were rinsed with 1 ml PBS and harvested with 500 ul of Trizol (Thermo Fisher, #15596018), according to manufacturer protocol. The recovered aqueous phase was additionally extracted once with 500 ul of Phenol:Chloroform:IAA, 25:24:1, pH 6.6 (Thermo Fisher, #AM9732) and once with 500 ul chloroform. The resulting aqueous phase was then ethanol-precipitated and resuspended in 30 ul 1x Turbo DNase buffer and treated with 2 ul Turbo DNase (Thermo Fisher, #AM1907) for 2 h at 37 °C. DNase was removed using the inactivation reagent from the same kit, according to manufacturer protocol.

RNA was reverse-transcribed using ProtoScript II First Strand cDNA Synthesis Kit (NEB, #E6560), according to manufacturer protocol, using the random primer mix and 4 ul RNA per 20 ul reaction. A duplicate reaction was included for one sample, substituting water for enzyme mix (a control for amplification of DNA). Reactions for qPCR were performed in a 96-well plate using iTaq Universal SYBR Green Supermix (Bio-Rad, #1725121) in a 20 ul volume with 1 ul template cDNA and 500 nM primer. SunTag primers were designed to amplify only intact (unspliced) reporter mRNAs (the primers bind and amplify a region that is spliced out of all cryptically spliced isoforms). Each cDNA sample (including the control for DNA amplification) was probed in triplicate using primers for both the SunTag reporter and the normalizing reporter. A series of 8 2-fold dilutions was generated from one of the samples and probed with each primer pair in duplicate for use as standard curves. qPCR was performed in a QuantStudio 6 Real-Time PCR System from Thermo Fisher.

To generate a standard curve for each primer pair, the dilution factor was plotted against the mean of the two duplicate CT values. These data were fit by an exponential curve, which was used to convert measured CT values to relative mRNA abundance. Triplicate samples were averaged and the result for the SunTag reporter divided by that of the normalizing reporter. The resulting ratio for each reporter was normalized to that of the no insert reporter. Each experiment was performed three times and the average and standard error of the mean reported.

<u>Determination of ZNF598 knock-down efficacy:</u>

To determine knock-down efficacy with siRNA treatment, U-2 OS cells stably expressing MCP-Halo-CAAX and scFV-sfGFP were seeded in a 6-well plate at a density of 90,000 cells per well. The following day, cells were transfected with a pool of 4 uM ZNF598-targeting siRNAs (Horizon, #L-007104-00-0005) or a pool of 4 uM non-targeting siRNA (Horizon, #D-001810-10-05) using RNAiMax (Thermo Fisher, #13778) according to manufacturer protocol. Approximately 24 h later, cells were transfected with a second dose of siRNAs. Approximately 4 h later, the media was changed to fresh DMEM + 10% FBS. Approximately 24 h later, media was aspirated, cells washed 1x with 2 ml PBS and scraped in 120 ul of a lysis buffer containing 50 mM Tris pH 8.0, 150 mM KCl, 1% triton X-100, 1 tablet EDTA-free protease inhibitor cocktail (Millipore Sigma, #11873580001), 20 units Turbo DNase (Thermo Fisher, #AM1907). Lysate was centrifuged for 5' at 21,000 g and the supernatant recovered. Total protein concentration in the lysate was determined using DC Protein Assay (Bio-Rad, #500-0114). Equal concentrations of total protein, along with a protein ladder (Bio-Rad, #161-0394) were loaded and resolved on 4-12% Criterion XT Bis-Tris protein gels (product info) and transferred to a PVDF membrane using Trans-Blot Turbo system (Bio-Rad). The membrane was blocked in TBST with 5% non-fat milk (Santa Cruz, #sc2325) for 1 h at room temperature. The gel was cut between the 50 and 75 kD marker bands, and the top half incubated overnight with a 1:5000 dilution of rabbit polyclonal ZNF598 primary antibody (Bethyl Laboratories, #A305-108A) in TBST with 5% non-fat milk at 4°C. The bottom half was incubated overnight in TBST with 5% non-fat milk at 4°C. Both membranes were washed 3x for 10 min with TBST. The top half of the blot was then incubated at room temperature for 45 min with an anti-rabbit HRP secondary antibody (Santa Cruz, #2357) at a 1:5000 dilution in TBST with 5% non-fat milk, while the bottom half was incubated at room temperature for 45 min with a beta-actin rabbit monoclonal antibody-HRP conjugate (Cell Signaling #5125) at a 1:5000 dilution in TBST with 5% non-fat milk. Both membranes were washed 3x for 10 min with TBST and developed using SuperSignal West Pico Plus ECL Substrate (Thermo Fisher, #34580). The top half of the membrane was supplemented with 10% SuperSignal West Femto Maximum Sensitivity Substrate (Thermo Fisher, #34095). Both membranes were exposed to Amersham Hyperfilm ECL (GE Healthcare, #28906839).

Harringtonine runoff experiments

U-2 OS cells stably expressing MCP-Halo-CAAX and scFV-sfGFP were seeded in 35 mm glass-bottom dishes (Cellvis #D35-20-1.5-N) at a density of 60,000 cells per dish. One or two days later, each dish was transfected with 800 ng reporter plasmid, using X-tremeGENE 9 DNA Transfection Reagent (Roche, #06365787001), at a ratio of 5 ul reagent to 1 ug DNA to 125 ul Opti-MEM Reduced Serum Medium (Thermo Fisher, #51985034). For experiments performed under ZNF598 depletion, cells were transfected the day after seeding with a pool of 4 uM ZNF598-targeting siRNAs (Horizon, #L-007104-00-0005) or a pool of 4 uM non-targeting siRNA (Horizon, #D-001810-10-05) using RNAiMax (Thermo Fisher, #13778) according to manufacturer protocol. Approximately 24 h later, cells were transfected with a second dose siRNAs. Approximately 4 h later, the media was changed to fresh DMEM + 10% FBS and cells were transfected with 800 ng reporter plasmid, using X-tremeGENE 9 DNA Transfection Reagent (Roche, #06365787001), at a ratio of 5 ul reagent to 1 ug DNA.

Approximately one day after transfection of the reporter, cells were dyed by adding JF-549 halo dye to dishes at a final concentration of 5 nM (*2*, *3*). 3-Indoleacetic acid (Sigma, #I2886) was added to the media at a final concentration of 0.5 mM either several hours post-transfection or at the time of dyeing. Approximately 1 h after adding dye, dishes were washed three times with DMEM + 10% FBS, returned to the incubator for approximately 30 min, and washed once more with DMEM + 10% FBS. Immediately prior to imaging, the media was switched to Leibovitz's L-15 media (Thermo Fisher, #11415) supplemented with 10% FBS. 3-Indoleacetic acid was maintained at a concentration of 0.5 mM throughout dyeing, washing and imaging.

During imaging, cells were maintained at 35-37°C. After identifying a cell to image, 0.75 ml media (1/3 total volume) was removed from the dish and harringtonine (Cayman Chemical,

#15361) was added to 9 ug/ml, mixed with a pipet and added back to the dish (3 ug/ml final concentration). Imaging was started 60 sec post-harringtonine addition in order to adjust the focus after mechanical perturbation of the microscope. Cells were excited sequentially with 488/561 nm lasers at 4% green/5% red power, respectively. The exposure time was 500 ms, frame rate one every 15 or 20 sec, and total imaging time 30 min.

Data were analyzed using a combination of custom MATLAB code, AirLocalize (*4*) and u-track (*5*). Particle detection was performed using AirLocalize, while u-track was used for tracking. Tracks shorter than 5 frames were discarded. A temporal overlap of at least five frames was required in order to link red and green tracks. mRNAs for which tracking was disrupted due to crossing paths with another mRNA were discarded.

To calculate the clearance time on each mRNA, we first determined when the green signal reached its maximum value, and then found the time after reaching the maximum value at which the signal fell below 10% of its maximum intensity. We performed the same analysis for the red channel. If loss of signal for red and green were coincident within three or fewer fames, we did not include the molecule for analysis, due to concerns about signal disappearance for reasons other than clearance of ribosomes (e.g. mRNAs leaving the membrane). Because the red signal was more difficult to track for the full 30 min than the green signal, we included molecules in the analysis for which green signal persisted after loss of red signal, as long as the green track was previously linked to a red track. mRNAs were not included if the mean signal of the first 4 frames was less than 10% of the maximum intensity (considered to be not translating at the start of the experiment).

Generation of stable cell lines

Lentiviral particles were generated by transfecting 293T cells with either MCP-Halo-CAAX or scFV-sfGFP plasmids along with viral packaging accessory plasmids. 48 hours following transfection, the viral supernatant was collected, spun down to remove cellular contents, and filtered through a 0.45 μm PVDF filter (Millipore SLHV013SL). The filtered supernatant was applied directly to U-2 OS cells (American Type Culture Collection HTB-96). Viral transduction was performed sequentially by first infecting U-2 OS cells with MCP-Halo-CAAX and performing fluorescence activated cell sorting (FACS) for positive cells. This positive population was then infected in the same manner with scFV-sfGFP and sorted for highly expressing cells.

Cell lines for live-cell imaging with stable expression of shRNAs for knock-down of ZNF598 were generated by lentiviral transduction of U-2 OS cells stably expressing MCP-Halo-CAAX and scFV-sfGFP using the psPAX2 packaging plasmid (Addgene #12260), pMD2.G envelope-expressing plasmid (Addgene #12259) and a lentiviral vector with either ZNF598-targeting shRNA (Sigma Aldrich Mission shRNA #TRCN0000073162) or mock shRNA (Addgene #1864). Transduced cells were selected by treatment with 5 ug/ml puromycin for one day.

Cell lines stably expressing the reporter mRNAs were generated by using the Flp-In method. U-2 OS cells containing a single Flp-In locus and stably expressing the T-Rex tet-On system (ThermoFisher) were a kind gift of Andrew Holland. Cells were first virally transduced with scFv-sfGFP as described above and sorted by FACS. The cells were then tranfected 1:1 mass ratio of pcDNA plasmid to pOG44 (ThermoFisher V600520) using X-tremeGENE 9 DNA Transfection Reagent (Roche, #06365787001) at a ratio of a ratio of 4 ul reagent to 1 ug DNA. Negative control plates received only pOG44. 48 hours following transfection, cells were trypsinized and re-plated in media containing 100 μg/mL hygromycin (InVivogen ant-hg-1) to begin positive selection. Individual colonies began to form ~1 week following transfection. Positive cells were pooled and expanded after complete cell death on the negative control plate.

smFISH probe labeling

smFISH probes were labeled following the protocol described in (*6*). Briefly, 20-mer plate DNA oligonucleotide probes ordered from IDT were pooled together and conjugated to amino-11-ddUTP (Lumiprobe A5040) at the 3'-end using terminal deoxynucleotidyl transferase (TdT) (Thermo Fisher EP0162). After purification by Spin-X centrifuge column (Corning 8161) with Bio Gel P-4 Beads (Bio Rad 1504124), the oligonucleotide-amino-11-ddUTP were labeled with Cy3-NHS ester (Lumiprobe 41020). After Cy3-lableing, the oligonucleotides were again purified to remove excessive dyes with Spin-X centrifuge column.

smFISH-immunofluorescence (smFISH-IF)

We performed smFISH-IF on U-2 OS cells stably expressing the mRNA reporters and scFV-sfGFP without MCP-Halo-CAAX. 18 mm #1 coverslips (Fisher 12-545-100) were base etched in 3M sodium hydroxide (Millepore Sigma 221465) for 30 minutes at room temperature. Coverslips were washed 4x with PBS (Corning 21-031-CV) and then coated for 1 hour at 37°C with 0.25 mg/mL rat tail collagen I (Gibco A1048301) diluted in 20 mM sodium acetate (Sigma-Aldrich S2889). The coverslips were washed 4x with PBS, and 40,000 cells were plated per well in DMEM supplemented with 10% FBS and pen/strep. 24 hours following plating, the media was exchanged and supplemented with 2 ug/mL doxycycline hyclate (Millipore Sigma, #D9891) and 250 uM 3-indole acetic acid (IAA) (Sigma-Aldrich I2886).

24 hours following induction smFISH-IF was performed as described (*7*). Briefly, all solutions were prepared in nuclease free water (Quality Biological 351-029-131CS). Cells were washed 3x with 1x PBS (Corning 46-013-CM) + 5 mM magnesium chloride (Sigma-Aldrich M2670-500G) (PBSM). Cell were then fixed for 10 minutes at room temperature in PBSM + 4% paraformaldehyde (Electron Microscopy Sciences 50-980-492). After fixation, samples were washed for 3x5 minutes in PBSM and then permeabilized in PBSM + 5 mg/mL BSA (VWR VWRV0332-25G ) + 0.1% Triton-X100 (Sigma-Aldrich T8787-100mL) for 10 minutes at room temperature. Cells were then washed 3x5 minutes in PBS and incubated for 30 minutes at room temperature in 2xSSC (Corning 46-020-CM), 10% formamide (Sigma-Aldrich F9037-100ML), and 5 mg/mL BSA (VWR VWRV0332-25G). After pre-hybridization incubation, cells were incubated for 3 hours at 37°C in 2xSSC (Corning 46-020-CM), 10% formamide (Sigma-Aldrich F9037-100ML), 1 mg/mL competitor *E. coli* tRNA (Sigma-Aldrich 10109541001), 10% w/v dextran sulfate (Sigma-Aldrich D8906-100G), 2 mM ribonucleoside vanadyl complex (NEB S1402S), 100 units/mL SUPERase In (Thermo Fisher AM2694), 60 nM SunTag_v4-Cy3 smFISH probes, and 1:1,000 chicken anti-GFP (Aves Labs GFP-1010). After incubation, the coverslips were washed 4x with 2xSSC (Corning 46-020-CM) + 10% formamide (Sigma-Aldrich F9037-100ML). The samples were then incubated with 2x20 minutes with a goat anti-chicken IgY secondary antibody conjugated to Alexa Fluor 488 (Thermo Fisher A-11039). The samples were then washed 3x with 2xSCC before being mounted on pre-cleaned frosted glass cover slides (Fisher 12-552-3) with ProLong Diamond antifade reagent with DAPI (Invitrogen P36962).

For smFISH-IF experiments performed after treatment with siRNAs, 90,000 U-2 OS cells expressing scFV-sfGFP and the corresponding reporter were seeded in a 6-well plate approximately 72 h prior to the experiment. Approximately 16-18 hours following seeding, cells were transfected with a pool of 4 uM or 21 uM ZNF598-targeting siRNAs (Horizon, #L-007104-00-0005), or a pool of 4 uM or 21 uM non-targeting siRNA (Horizon, #D-001810-10-05) using RNAiMax (Thermo Fisher, #13778) according to manufacturer's protocol. 6 hours following the initial transfection, cells were re-plated onto coverslips as described above. 24 hours prior to the smFISH-IF protocol, cells were transfected with a second dose of siRNA and the media was supplemented with 2 ug/mL doxycycline hyclate (Millipore Sigma, #D9891) and 250 uM 3-indole acetic acid (IAA) (Sigma-Aldrich I2886). From this point, the smFISH-IF protocol was performed as described above.

Microscope

Live cell data were acquired on a custom inverted wide-field Nikon Eclipse Ti-E microscope equipped with three Andor iXon DU897 EMCCD cameras (512x512 pixels), Apochromatic TIRF 100x Oil Immersion Objective Lens/1.49 NA (Nikon MRD01991), linear encoded Stage XY-stage with 150 micron Piezo Z (Applied Scientific Instrumentation), and LU-n4 four laser unit with solid state 405 nm, 488 nm, 561 nm, and 640 nm lasers (Nikon), a TRF89901-EM ET-405/488/561/640nm Laser Quad Band Filter Set for TIRF applications (Chroma), and Nikon H-TIRF system. The x-y pixel size was 160 nm.

Fixed cell data were acquired on a custom wide-field inverted Nikon Ti-2 wide-field microscope equipped with 60x 1.4NA oil immersion objective lens (Nikon), Spectra X LED light engine (Lumencor), and Orca 4.0 v2 scMOS camera (Hamamatsu). The x-y pixel size was 107.5 nm and the z-step size was 300 nm. Both microscopes were under the automated control of the Nikon Elements software.

Long-term imaging of steady-state translation

U-2 OS cells stably expressing MCP-Halo-CAAX and scFV-sfGFP were seeded in 35 mm glass-bottom dishes (Cellvis #D35-20-1.5-N) at a density of 60,000 cells per dish. One day later, each dish was transfected with 400 ng reporter plasmid, using X-tremeGENE HP DNA Transfection Reagent (Roche, 6366236001), at a ratio of 4 ul reagent to 1 ug DNA to 100 ul DMEM (Corning, 10-013-CV). Approximately one day after transfection of the reporter, cells were dyed by adding JF-549X halo dye (a gift from Dr. Luke Lavis) to dishes at a final concentration of 10 nM (*2*, *3*). Approximately 1 h after adding dye, dishes were washed three times with DMEM + 10% FBS, and media was switched to FluoroBrite DMEM (Thermo Fisher #A1896701) + 10% FBS. 3-Indoleacetic acid (Sigma, #I2886) was maintained at a concentration of 0.5 mM throughout dyeing, washing and imaging. During imaging, cells were maintained at 37°C and 5% $CO_2$. Cells were excited simultaneously with 488/561 nm lasers at 4% green/6% red power, respectively. The exposure time was 500 ms, frame rate one every 15 sec, and total imaging time 3 h.

Data were analyzed using a combination of custom MATLAB code, AirLocalize (*4*) and u-track (*5*). Particle detection was performed using AirLocalize, while u-track was used for tracking. Tracks shorter than 5 frames were discarded. A temporal overlap of at least five frames was required in order to link red and green tracks. mRNAs for which tracking was disrupted due to crossing paths with another mRNA were discarded.

Oxford Nanopore sequencing

Reporters were expressed in HEK293T cells to maximize transfection efficiency and thus sequencing depth of the reporters. This is important since the vast majority of sequencing reads map to endogenous mRNAs, limiting the ability to acquire reads that map to the reporter. 1,500,000 HEK293T cells were seeded in 10 cm dishes, and transfected the following day with 17.5 ug of pcDNA_CMV_ST or pcDNA_CMV_ST_A60, 45 ul Lipofectamine 3000 reagent (Thermo Fisher, #L3000008), 35 ul P3000 reagent and 1.5 ml Opti-MEM Reduced Serum Medium (Thermo Fisher, #51985034). Two days later, cells were washed with 5 ml PBS and harvested in 1 ml Trizol (Thermo Fisher, #15596018), according to manufacturer protocol. The recovered aqueous phase was additionally extracted once with 500 ul of Phenol:Chloroform:IAA, 25:24:1, pH 6.6 (Thermo Fisher, #AM9732) and once with 500 ul chloroform. The resulting aqueous phase was then ethanol-precipitated and resuspended in 50 ul dd$H_2O$. 5.5 ul of 10x Turbo DNase buffer and 2 ul Turbo DNase (Thermo Fisher, #AM1907) were added to each sample and reactions incubated for 10 min at 37 °C. DNase was removed using the inactivation reagent from the same kit, according to manufacturer protocol. The total RNA sample was polyA-selected using NEBNext Poly(A) mRNA Magnetic Isoluation Module (NEB, #E7490), according to manufacturer protocol with two modifications: We used three times the amount of beads recommended and eluted in a 15 ul volume. Library prep was performed using the Nanopore Direct RNA Sequencing Kit (Oxford Nanopore #SQK-RNA002) and sequenced at the Johns Hopkins University Genetic Resources Core Facility on a GridION instrument.

Reads were mapped to reference plasmid sequences using the minimap2 program (*8*). Splicing status was assessed using custom code written in python with the pysam module. The common splice acceptor site for the four spliced isoforms is located at position 2921 of the reference sequence. To estimate the percentage of reads representing each isoform, it was necessary to count the total number of reads spanning enough sequence surrounding the splice junction such that it was possible to assess splicing status. Thus, we counted the number of reads fully spanning the region from position 2870 to 2972 (874 reads for pcDNA_CMV_ST, 463 reads for pcDNA_CMV_ST_A60) and divided the count for each isoform by that number.

### Supplementary Text

#### Calculating theoretical brightness of cryptically spliced SunTag isoforms

To determine the relative brightness of cryptically spliced SunTag isoforms, we calculated the brightness of each translation site as $F(t) = \theta(\sum_{i=1}^{N} i p_i(t) + N \sum_{i=N+1}^{M} p_i(t))$, where $\theta$ is the brightness of a single SunTag-labeled epitope, N is the number of SunTag epitopes in the reporter (N = 23 for intact reporter), M is the total number of segments of SunTag length in the reporter (M = 52 for intact reporter), and $p_i(t)$ is the probability that a ribosome is located at segment $i$ at time $t$ (*1*). We assumed that $p_i(t)$ is constant for all $i$. For each splice isoform, we determined the number of SunTag epitopes (N) and the total length of the open reading frame (M) to calculate the brightness relative to the intact reporter.

#### Calculating the number of translating ribosomes from FISH-IF data

All fixed cell imaging analysis was performed with existing and custom software packages in MATLAB and as previously described (*7*). Spot detection of both smFISH and immunofluorescence channels was performed independently using FISH-Quant (*9*). FISH-Quant uses Gaussian fitting to determine sub-pixel spot localizations and integrated spot intensities. In the mRNA channel, only single transcripts in the cytoplasm were considered. After determining mRNA localizations, FISH-Quant's transcription site fitting algorithm was used to quantify the integrated intensity of the translation site. In brief, using the mRNA localization, a 11x11 pixel bounding box was drawn around each mRNA and designated as potential translation site. Single mature polypeptides were detected across the entire image outside of the potential translation sites.

These single peptides were thresholded based on their Gaussian fitting parameters (width and intensity) and inspected to ensure accuracy. These single peptides were averaged using FISH-Quant to generate an idealized point-spread-function to calculate the integrated intensity for a single peptide. The potential translation sites were fit to a Gaussian centered on the brightest pixel within the boundingbox. The fitting results were again filtered based on shape, intensity, and distance from the original mRNA positions. Failure to converge on an accurate fit based on these parameters resulted in an integrated intensity of 0. The integrated intensity of the single particle was used to calculate the number of nascent peptides within the translation site. Because not all ribosomes on the mRNA have translated the full SunTag sequence, we imposed a correction factor to determine the number of ribosomes on the mRNA from the raw number of nascent peptides. The correction is based on the relative proportion of the open reading frame that contains the SunTag sequence:

$$\#\, Ribosomes = \frac{N}{N - n/2} \times \frac{P}{I_{Single}}$$

where N is the total length of the protein in either nucleotides or peptides, n is the SunTag length, P is the intensity of the translation site and $I_{Single}$ is the mean intensity of the single protein distribution(*7*). The correction factor ($\frac{N}{N-n/2}$) for each open reading frame is listed in the table below.

| Reporter | Nascent Peptide Correction Factor |
|---|---|
| no insert | 1.309660574 |
| poly(A)36 | 1.305827746 |
| poly(A)60 | 1.303324808 |

These correction factors assume a uniform density of ribosomes along the mRNA. Because we observed increased ribosome occupancy on poly(A) reporters caused by ribosome queuing, ribosome density on these mRNAs is likely biased towards the 3'-end of the mRNA. Thus, the correction factors listed in the table may result in slight overestimation of the ribosome load on poly(A) reporters. However, even if we assume the extreme case in which all ribosomes on the poly(A)60 reporter have synthesized the full SunTag array—and thus we do not apply the correction—the poly(A)60 reporter is still occupied by an average of ~45 ribosomes.

Calculating the frequency of foci separation during long-term imaging

To calculate the frequency of foci separation, we first determined the total length of time that each mRNA was observed in a translating state. The sum of single mRNA translation times over all molecules and all cells within a given sample is the total observation time for that sample. We then counted the number of times that foci separation was observed for all molecules and all cells within the sample and divided the foci separation count by the total observation time to get the foci separation frequency.

Testing statistical significance of difference between distributions of ribosome number per mRNA:

To test distributions of number of ribosomes per mRNA for statistically significant differences (Figures 2B, 4C and 4D), distributions were first transformed by taking the square root of all values to reduce skewness. The two two-sample t-test was then applied to the transformed distributions.

A.

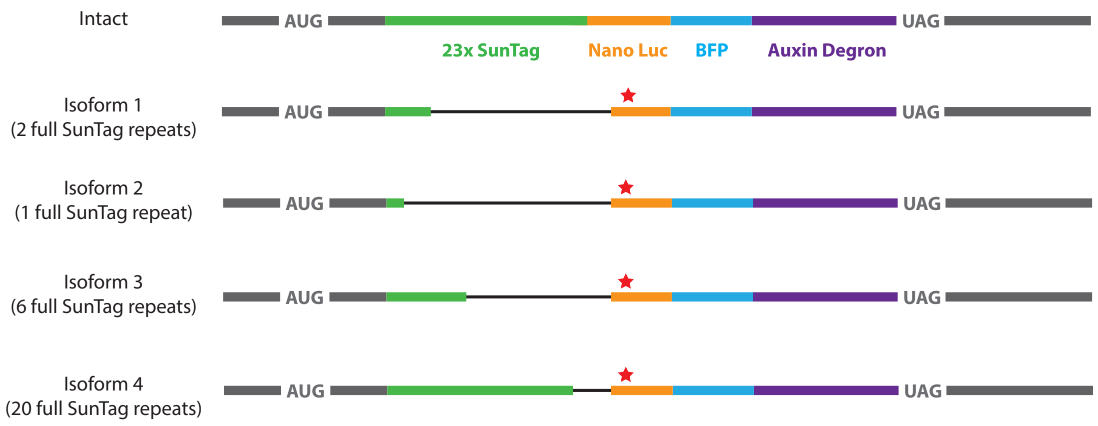


B.

| | % of reads for no insert reporter | % of reads for polyA(60) reporter | Calculated relative SunTag intensity |
|---|---|---|---|
| Expected (intact) | 73.2 | 69.6 | 1 |
| Isoform 1 | 16.9 | 19.1 | 0.007 |
| Isoform 2 | 6.1 | 7.1 | 0.002 |
| Isoform 3 | 2.1 | 1.9 | 0.031 |
| Isoform 4 | 1.7 | 2.3 | 0.257 |

**Fig. S1.**

Direct sequencing of mRNA reveals cryptic splicing in the SunTag region of the reporters. Reporter plasmids were transfected into HEK293T cells and mRNA was sequenced by Oxford Nanopore direct RNA sequencing. (A) Cartoon schematic of the intact reporter, shown for reference, and the four detected splice isoforms. The thin black line indicates the intronic region. All isoforms share the same splice acceptor site and result in deletion of the first 33 codons of Nano Luciferase. Additionally, all splice events cause the same shift in frame, resulting in a stop codon 50 bp downstream of the splice junction (indicated by red star). Thus, none of these isoforms contribute to Nano Luciferase signal. (B) The prevalence of each isoform out of the total pool of reads with fragment starts and ends at least 50 bp upstream and downstream of the splice acceptor site, and the theoretical intensity based on the number of SunTag repeats and the ORF length (see supplemental text). The frequency of each isoform is not affected by the presence of poly(A). Isoforms 1 and 2 are not detected, as their calculated intensity is less than 1% that of the intact reporter. Isoforms 3 and 4 can in principle be detected; however together they account for only 4% of total reads and their abundance is the same in both samples.

A.

B.

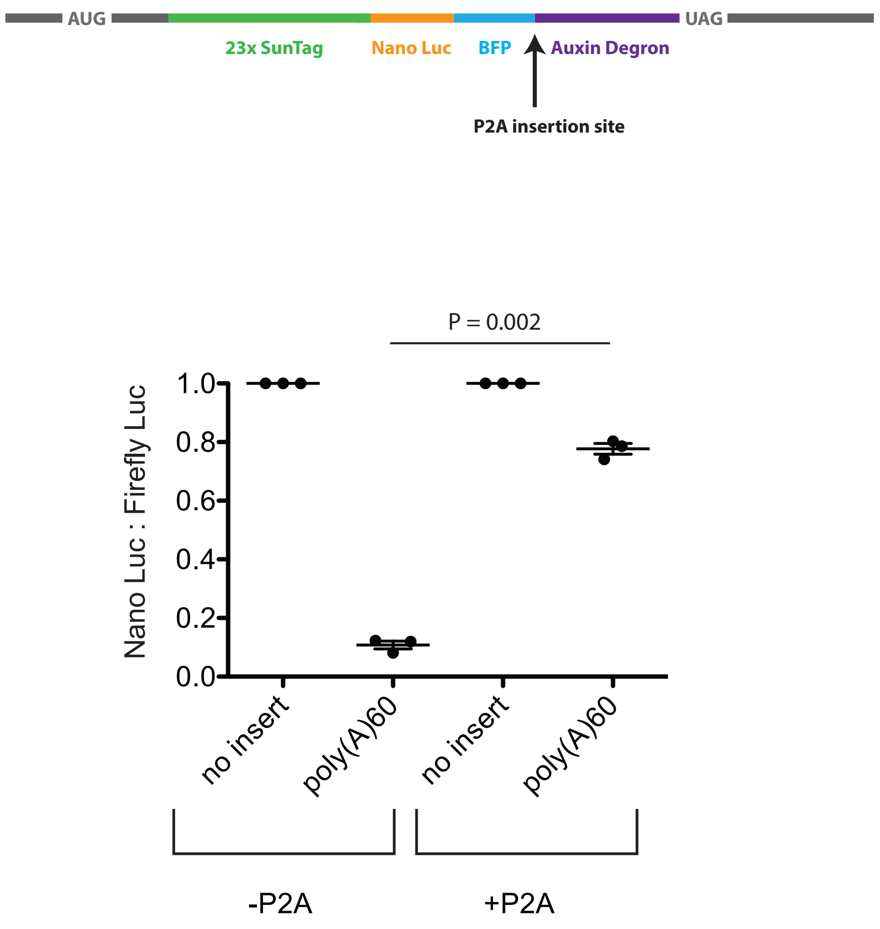


**Fig. S2.**

Release of Nano Luciferase from the ribosome prior to stalling at the poly(A) site restores protein output. (A) A P2A sequence was inserted just upstream of the Auxin Degron, resulting in peptide release 705 nt upstream of the beginning of the poly(A) insertion site. (B) Luciferase assay to measure protein output in the presence or absence of the P2A sequence. Nano Luciferase signal is normalized to Firefly Luciferase signal from a co-transfected plasmid and the resulting ratios are normalized to the no insert reporter within each P2A context. Error bars represent the sem of 3 independent experiments. p-values calculated by paired-sample t-test.

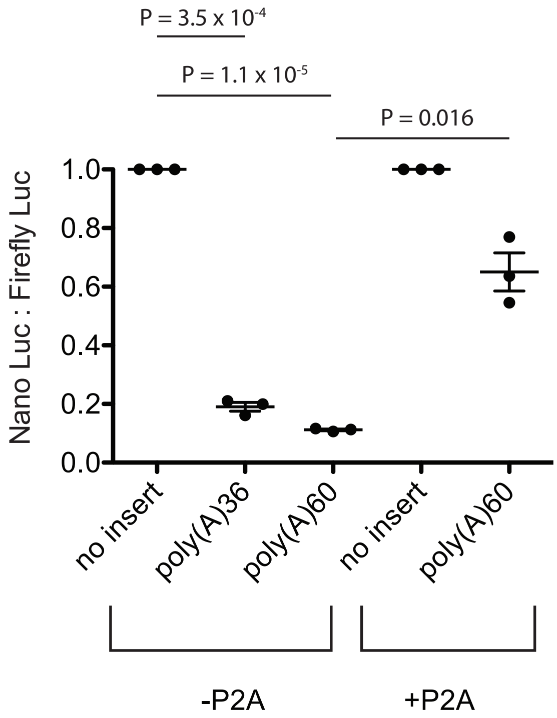


**Fig. S3.**
Luciferase assay performed in HEK293T cells. Nano Luciferase signal is normalized to Firefly Luciferase signal from a co-transfected plasmid and the resulting ratios are normalized to the no insert reporter. Error bars represent the sem of 3 independent experiments. p-values calculated by paired-sample t-test.

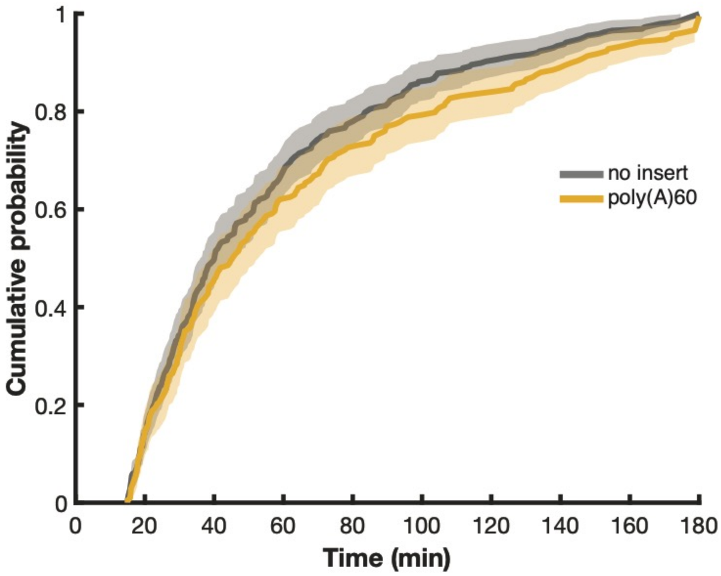


**Fig. S4.**
Cumulative probability distribution of mRNA observation times. For no insert, median observation time is 40.3 min; for poly(A)60, median observation time is 45.5 min. Only mRNAs observed for at least 15 min were included in the analysis. For no insert: 6 cells, 287 mRNA. For poly(A)60: 5 cells, 208 mRNA.

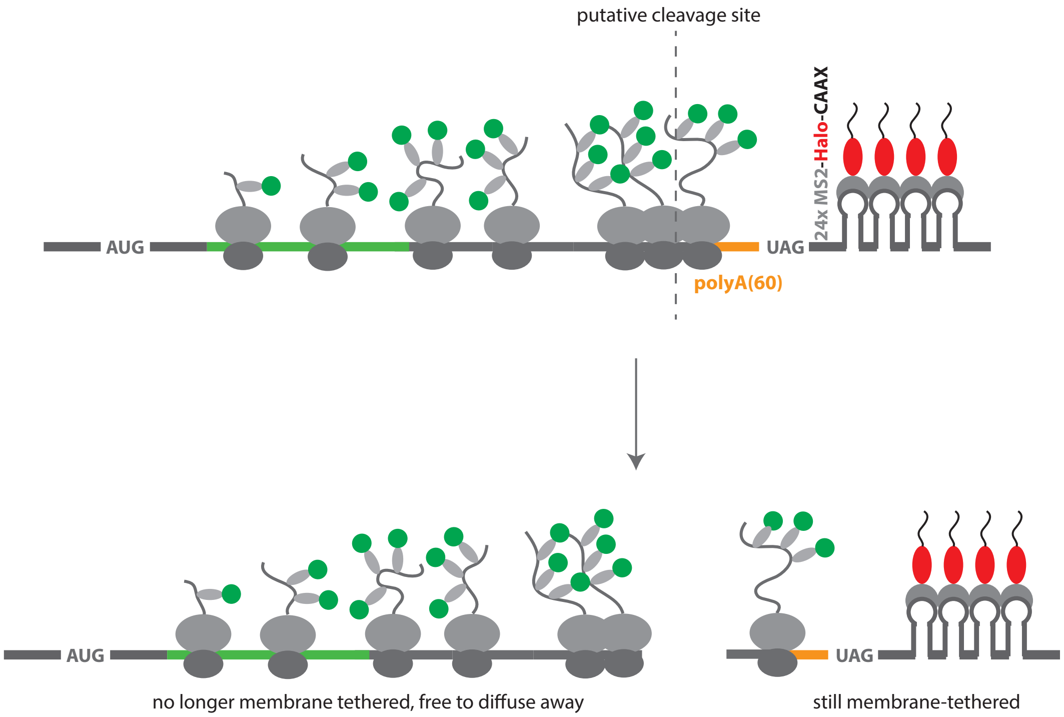


**Fig. S5.**

Cartoon showing the expected mRNA cleavage location based on previous work in yeast (*10*). If the mRNA is cleaved in the A site of the first collided ribosome, all ribosomes on the 5’ mRNA fragment are severed from the stem loops that tether the mRNA to the membrane. Thus, the majority of the green signal should diffuse away from the tethered red signal (foci separation).

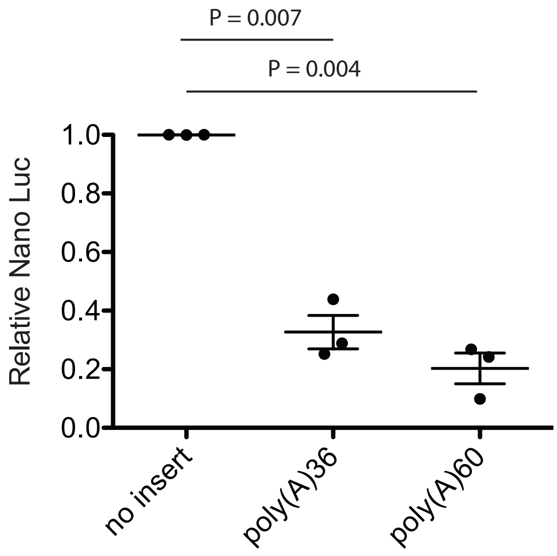


**Fig. S6.**

Luciferase assay to measure protein output in cell lines stably expressing the SunTag reporters. Equal numbers of cells were seeded in wells of a 96-well plate and induced with 2 ug/ml doxycycline approximately 24 h prior to measurement. Nano Luciferase signal for all three reporters was normalized to the signal for the no insert reporter. Error bars represent the sem of 3 independent experiments. p-values calculated by paired-sample t-test.

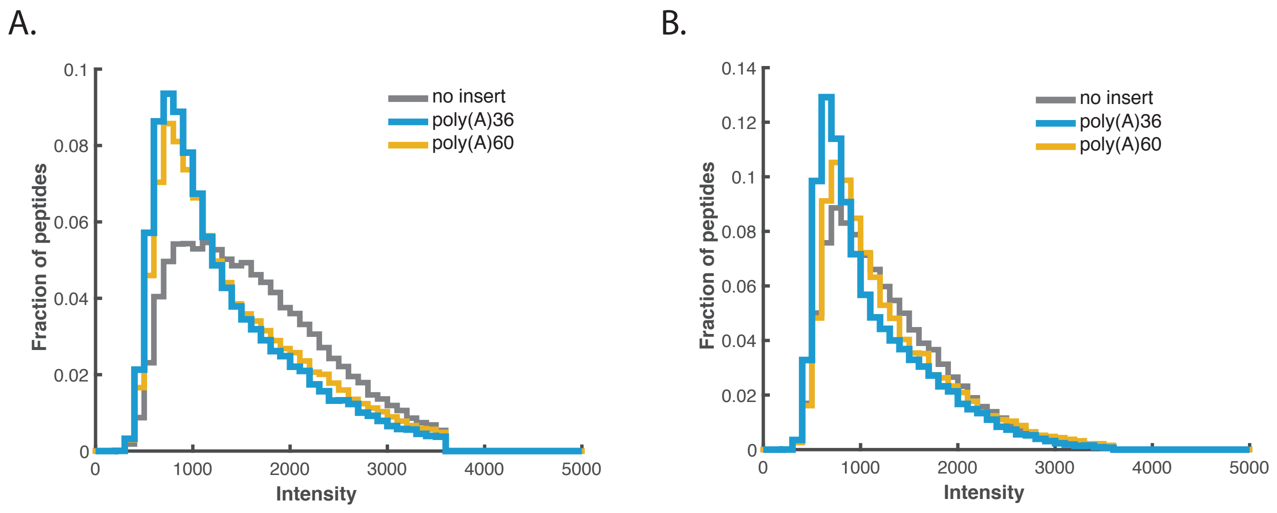


**Fig. S7.**

Raw intensity histogram of detected fully synthesized single peptides from two independent experiments. Number of ribosomes (Figure 2C) was calculated by dividing the SunTag intensity of each mRNA by the mean intensity of the fully synthesized single peptides detected from the respective sample. The number of ribosomes (Figure 2C) reflects combined data from both experiments.

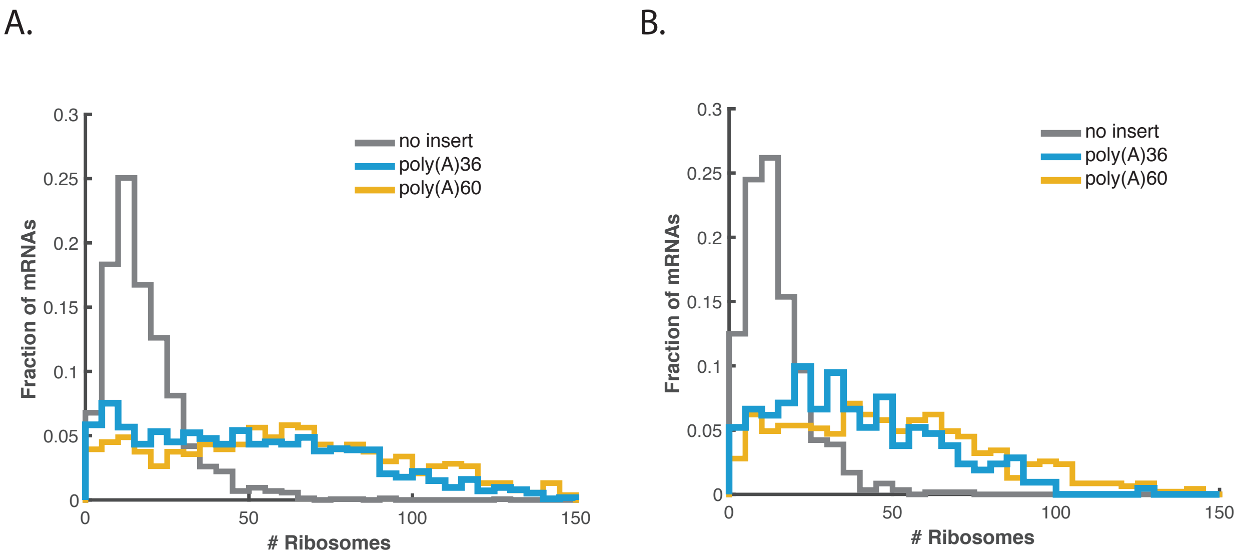


**Fig. S8.**

Two independent replicate smFISH-IF experiments were performed for the no insert, poly(A)36 and poly(A)60-expressing cell lines. The data shown in main figure 2C reflect the aggregate data from both replicates, which are plotted separately here. A) Replicate 1 B) Replicate 2.

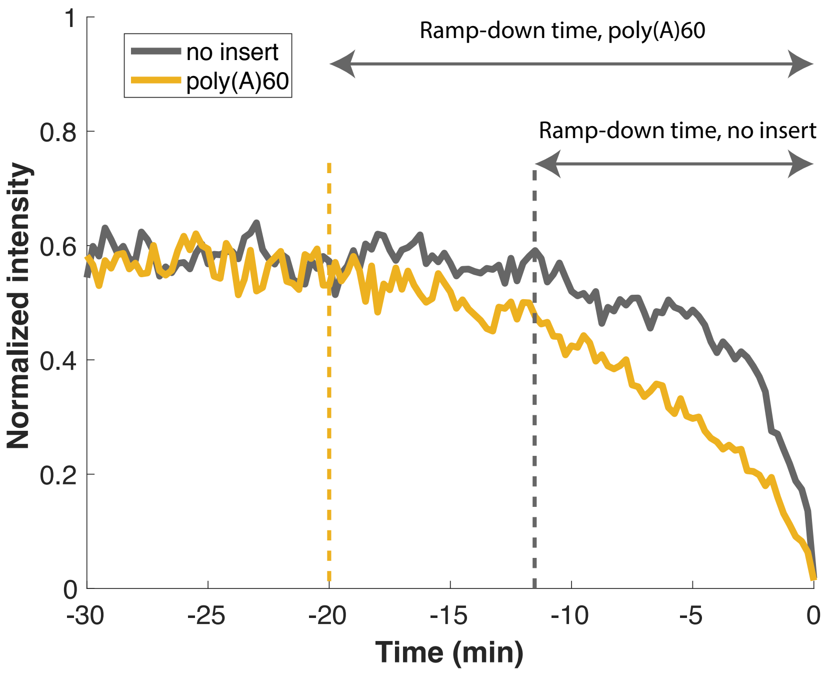


**Fig. S9.**

Translation shut-down proceeds more slowly on poly(A) mRNAs than on no insert mRNAs. The plot depicts the average normalized intensity of translation ramp-downs observed during live-cell imaging. For each instance of a ramp, intensity was measured starting 30 min before complete shut-down until complete shut-down, and fluorescence was normalized to the maximum intensity of the trace. The average of all normalized traces is plotted. Dotted vertical lines indicate the approximate time at which the intensity starts to decline, and double arrows indicate the approximate total time between when the intensity starts to decline and total translation shut-down. Note that ramp-down events are only included if they exhibit steady-state translation for at least 30 min prior to complete shut-down. For no insert, N = 21 traces; for poly(A)60, N = 26 traces.

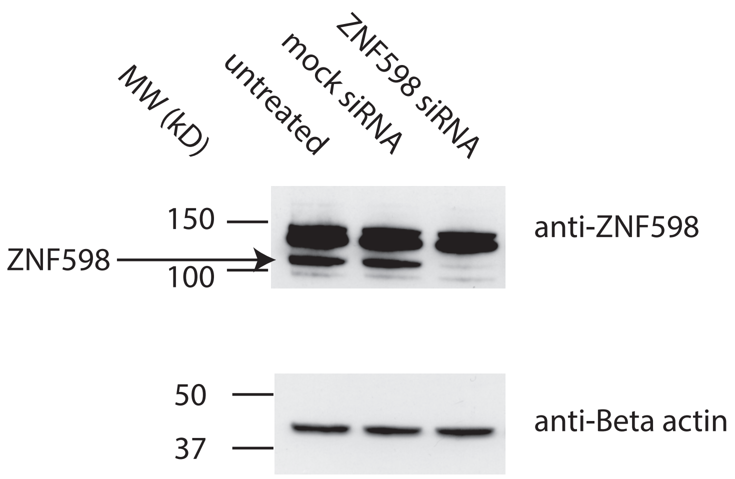


**Fig. S10.**
ZNF598 is effectively depleted by treatment with 4 nM siRNAs.

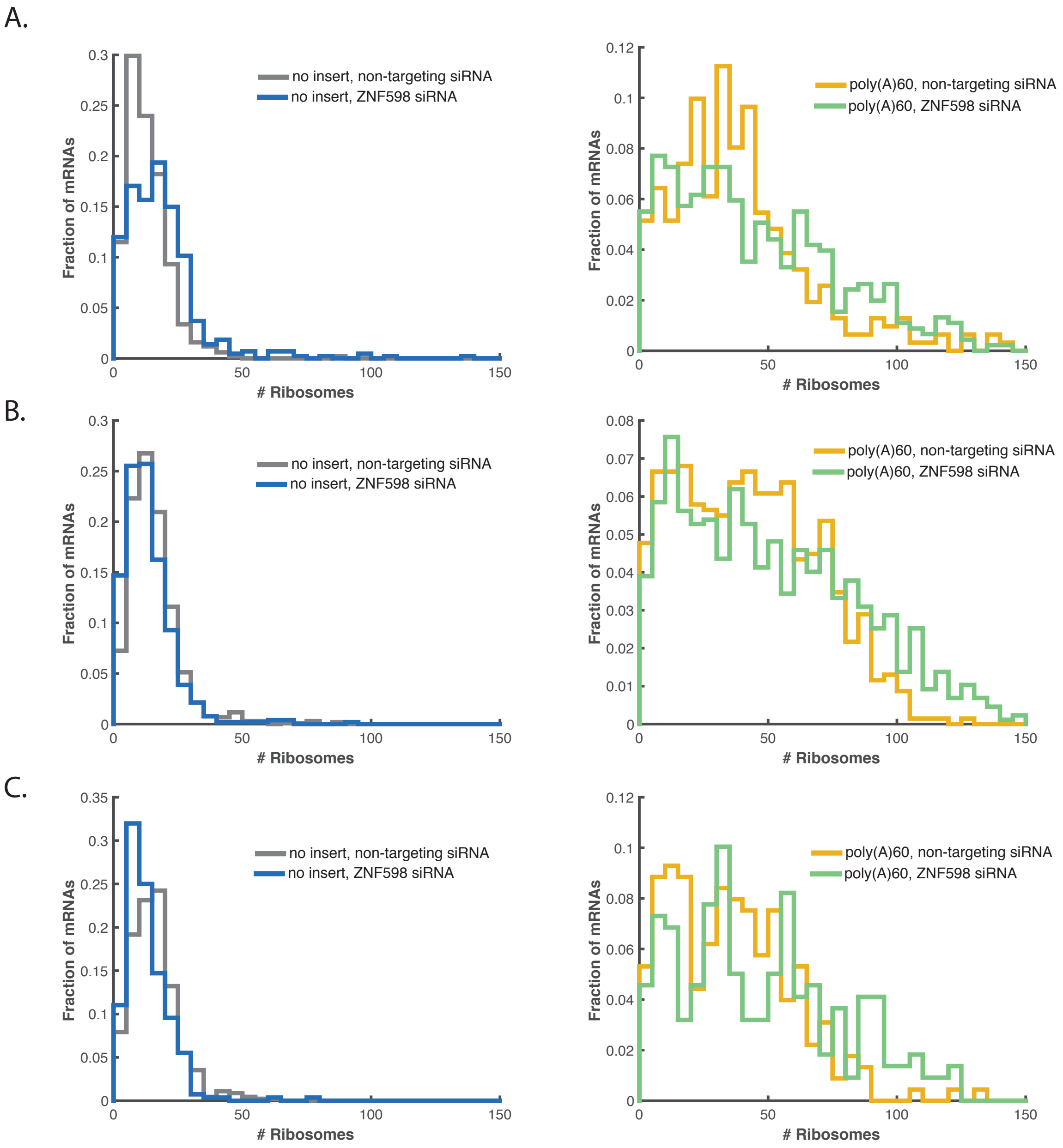


**Fig. S11.**

Three independent FISH-IF experiments were performed on siRNA-treated cells. (A) 1st replicate at 4 nM siRNA. Left: no insert reporter; Right: poly(A)60 reporter. (B) 2nd replicate at 4 nM siRNA. Left: no insert reporter; Right: poly(A)60 reporter. (C) This experiment was performed at 21 nM siRNA. The data shown in main figures 4C and 4D is the aggregate of the two replicate experiments at 4 nM siRNA.

**Movie S1.**

Imaging of translation on the no insert SunTag reporter. Top panel: cropped area of a cell, with one featured mRNA indicated with a green circle. Although imaging was performed for 3 h, the movie spans only 2 h because the featured mRNA was not tracked for the entire experiment. Red channel: mRNA; green channel: SunTag. Bottom pannel: Fluorescence intensity over time plotted for the featured molecule.

**Movie S2.**

Imaging of translation on the poly(A)60 SunTag reporter. Top panel: cropped area of a cell, with one featured mRNA indicated with a green circle. Although imaging was performed for 3 h, the movie spans only 2 h because the featured mRNA was not tracked for the entire experiment. Red channel: mRNA; green channel: SunTag. Bottom pannel: Fluorescence intensity over time plotted for the featured molecule.

**Movie S3.**

Example of red-green foci separation on the poly(A)60 SunTag reporter. Top panel: cropped area of a cell, with one featured mRNA indicated with a green circle. Red channel: mRNA; green channel: SunTag. Bottom pannel: Fluorescence intensity over time plotted for the featured molecule. Foci separation occurs after ~35 min.

**Movie S4.**

Harringtonine runoff experiment with a cell expressing the no insert reporter. Top panel: cropped area of a cell, with one featured mRNA indicated with a green circle. Red channel: mRNA; green channel: SunTag. Bottom pannel: Fluorescence intensity over time plotted for the featured molecule.

**Movie S5.**

Harringtonine runoff experiment with a cell expressing the poly(A)36 reporter. Top panel: cropped area of a cell, with one featured mRNA indicated with a green circle. Red channel: mRNA; green channel: SunTag. Bottom pannel: Fluorescence intensity over time plotted for the featured molecule.

**Movie S6.**

Harringtonine runoff experiment with a cell expressing the poly(A)60 reporter. Top panel: cropped area of a cell, with one featured mRNA indicated with a green circle. Red channel: mRNA; green channel: SunTag. Bottom pannel: Fluorescence intensity over time plotted for the featured molecule.

**Movie S7.**

Harringtonine runoff experiment with a cell expressing the no insert reporter in mock siRNA-treated cells. Top panel: cropped area of a cell, with one featured mRNA indicated with a green circle. Red channel: mRNA; green channel: SunTag. Bottom pannel: Fluorescence intensity over time plotted for the featured molecule.

**Movie S8.**

Harringtonine runoff experiment with a cell expressing the poly(A)60 reporter in mock siRNA-treated cells. Top panel: cropped area of a cell, with one featured mRNA indicated with a green circle. Red channel: mRNA; green channel: SunTag. Bottom pannel: Fluorescence intensity over time plotted for the featured molecule.

**Movie S9.**

Harringtonine runoff experiment with a cell expressing the no insert reporter in ZNF598 siRNA-treated cells. Top panel: cropped area of a cell, with one featured mRNA indicated with a green circle. Red channel: mRNA; green channel: SunTag. Bottom pannel: Fluorescence intensity over time plotted for the featured molecule.

**Movie S10.**

Harringtonine runoff experiment with a cell expressing the poly(A)60 reporter in ZNF598 siRNA-treated cells. Top panel: cropped area of a cell, with one featured mRNA indicated with a green circle. Red channel: mRNA; green channel: SunTag. Bottom pannel: Fluorescence intensity over time plotted for the featured molecule.
